# Spatiotemporal dynamics and assembly process differ in fungal communities across contiguous habitats in tropical forests

**DOI:** 10.1101/2024.09.20.614103

**Authors:** Chieh-Ping Lin, Yu-Fei Lin, Yu-Ching Liu, Mei-Yeh Jade Lu, Huei-Mien Ke, Isheng Jason Tsai

## Abstract

**Background:** The variation in fungal community composition within a single habitat space has been extensively studied in forest ecosystems. However, the spatial and temporal distribution of fungi across contiguous habitats, particularly at a local scale and in tropical regions, remains underexplored. In this study, we examined the fungal community composition across multiple habitats proximal to each other over two seasons in seven Fagaceae species in Taiwanese broadleaf forests. We tested how local spatial scale and habitat influence community assembly.

**Results:** Using a metabarcoding approach, we sequenced ITS3/ITS4 amplicons from 864 samples collected from four distinct habitats—leaves, twigs, litter, and soil. We identified 11,600 fungal amplicon sequence variants (ASVs), with community composition differing significantly between habitats proximal to each other. Phyllosphere (leaves and twigs) fungi exhibited higher diversity compared to soil. Habitat type and long-term precipitation emerged as the most influential factors driving fungal diversity and composition, with a clear distance-decay relationship observed in leaf and twig but not in soil. Random forest analysis accurately classified habitats based on ASVs’ relative abundances, with strong predictors were mostly endemic ASVs prevalent in soil. Misclassified samples were due to secondary contact of fungi between adjacent habitats. Co-occurrence network analysis revealed more complex and deterministic networks in leaf and twig habitats, while soil was driven by stochastic processes and contained most endemic ASVs. A *Cladosporium* sp. emerged as a keystone species, maintaining network stability across forests.

**Conclusion:** This study reveals how local spatial variation and habitat shape distinct fungal communities in tropical forests, with deterministic processes dominating in some habitats and stochasticity playing a key role in others. We show local fungal taxa were strong habitat predictors and drivers of community cohesion. These findings highlight the importance of studying coexisting habitats to gain a deeper understanding of fungal biogeography and ecosystem function.

## Background

Forests are highly heterogeneous ecosystems, comprising a diverse array of distinct habitats. Even at the scale of individual trees, spatial heterogeneity is evident with various compartments ranging from fresh leaves to soil, each supporting distinct fungal communities. These fungi provide niche-specific processes as well as interface-mediated responses, which collectively contribute to the health and functioning of the forest ecosystem [1–6]. For example, phyllosphere fungi play essential roles in nutrient cycling and organic matter decomposition following the fall of leaves, transforming materials into absorbable forms for plants [7, 8]. While numerous studies have explored fungal diversity within specific habitats on a global scale [9, 10], investigating the spatiotemporal patterns of fungal communities across multiple proximal habitats at a local scale can reveal how these communities assemble, maintain, or partition themselves. Such an approach can yield crucial insights into the biogeography of microorganisms and ecosystem processes [11, 12].

Spatiotemporal variability of fungal communities has been extensively studied in other systems, particularly in soil environments where mycorrhizal fungi play key roles [11]. A nearly universal biogeographical pattern observed is the decay in community similarity with spatial distance [13, 14]. This distance-decay relationship can result from dispersal limitation, whereby the dispersal of species tends to decrease as the physical distance from the source increases [15]. Additionally, geographically proximate locations often exhibit comparable environmental conditions, which serve to reinforce species similarity within these regions. Furthermore, the impact of these drivers on fungal communities varies across different habitats, leading to distinct patterns of community assembly [16]. These effects are inherently dependent on the spatial scale of the investigation [17]. At smaller spatial scale, stochastic processes may be dominant where environments are more homogenous. However, as spatial distance increases, environmental heterogeneity also increases (e.g., variations in vegetation, soil and litter chemistry, and climate), and community composition may then be more strongly determined with this heterogeneity [18]. Determining whether community assembly is driven by neutral processes or deterministic factors, such as niche differentiation, is crucial to understand the spatiotemporal dynamics and ecological drivers of fungal communities.

Fungal community composition can vary significantly across different microhabitats of forest foreground, such as leaves, twigs, topsoil, and litter, even when these microhabitats are in close proximity. These microhabitats differed extensively in environmental factors such as chemical nutrient availability, but at the same time were connected via co-inhabiting fungi [19] and microbial-mediated reactive interfaces [20]. Litter, for instance, can be particularly dynamic and experiences substantial seasonal changes over very small distances [21], which can act as bottleneck events due to the influx of new litter [12]. The litter-soil interface is essential for nutrient cycling and organic matter decomposition, impacting soil fertility and plant health. Examining the fungal community of the forest ecosystem as a whole would enable us to quantify the diversity differences between different environments that were otherwise biased by methodologies [13] and to investigate the relationship between these environments and delineate ecosystem processes that affect single or multiple habitats simultaneously [19].

Tropical and subtropical forests, which cover 56% of the global forest area and support 42.8% of the world’s tree species, are among the most biodiverse ecosystems [19, 22]. Taiwan is a continental island with 60.7% of its land area covered by forests [23], which is more than double the global average of 30.2% and it ranks as the fifth highest nation in terms of tree density [24]. In particular, the broadleaf forests are home to a diverse range of Fagaceae plant and microbial species [25], yet studies on the mycobiome of Fagaceae species remain limited. Oak trees, which are keystone species in many ecosystems, play a crucial role in maintaining forest structure and function [26–28]. Despite the ecological importance of these trees, there has been a paucity of research on the fungal communities associated with them. Given Taiwan’s rich biodiversity, varied environmental conditions, and designation as a priority area for fungal conservation [29], it serves as an ideal place for studying fungal communities. Understanding the mycobiome of Fagaceae species in Taiwanese forests can provide valuable insights into the ecological processes governing these ecosystems and contribute to more effective forest management and conservation strategies.

In this study, we investigated fungal diversity across closely situated microhabitats—leaves, twigs, soil, and litter— associated with seven Fagaceae species in two broadleaf forests and a botanical garden in Taiwan to understand how these different environments influence the assembly and variability of fungal communities at a very local scale. Using a metabarcoding approach, we sequenced ITS3/ITS4 amplicons from 864 samples collected across four different habitats: leaves, twigs, litter, and topsoil from the same trees. By examining the spatiotemporal variation in mycobiomes at two time points, we sought to determine the influence of factors such as spatial distances, habitat type, season, and host species on fungal community composition. Our comprehensive analysis highlights the dynamic nature of fungal communities in the foreground of Taiwanese Fagaceae forests and underscores the significant role of environmental factors in shaping these communities. This study provides valuable insights into the complexity and variability of fungal diversity in these ecosystems, paving the way for future research on microbial biogeography and forest ecosystems as a whole.

## Methods

### Sample collection

To investigate the spatiotemporal variations in the forest mycobiomes, we collected samples from leaves, twigs, litter and topsoil of 38 trees belonging to seven Fagaceae species (*Quercus stenophylloides* n=14, *Quercus glauca* n=7, *Quercus morii* n=2, *Quercus pachyloma* n=3, *Castanopsis fargesii* n=2, *Lithocarpus hancei* n=5, *Lithocarpus glaber* n=5) at two locations in Nantou County (Puli Township and Ren’ai Township) and at the Fushan Botanical Garden in Yilan County, Taiwan (**Fig. 1a and 1b**). Samples were collected at two time points per location, with Nantou samples harvested in April and October 2022, and Fushan samples in July and December 2022. The Fushan samples were designated as artificial Fagaceae woodlands.

**Fig. 1.**
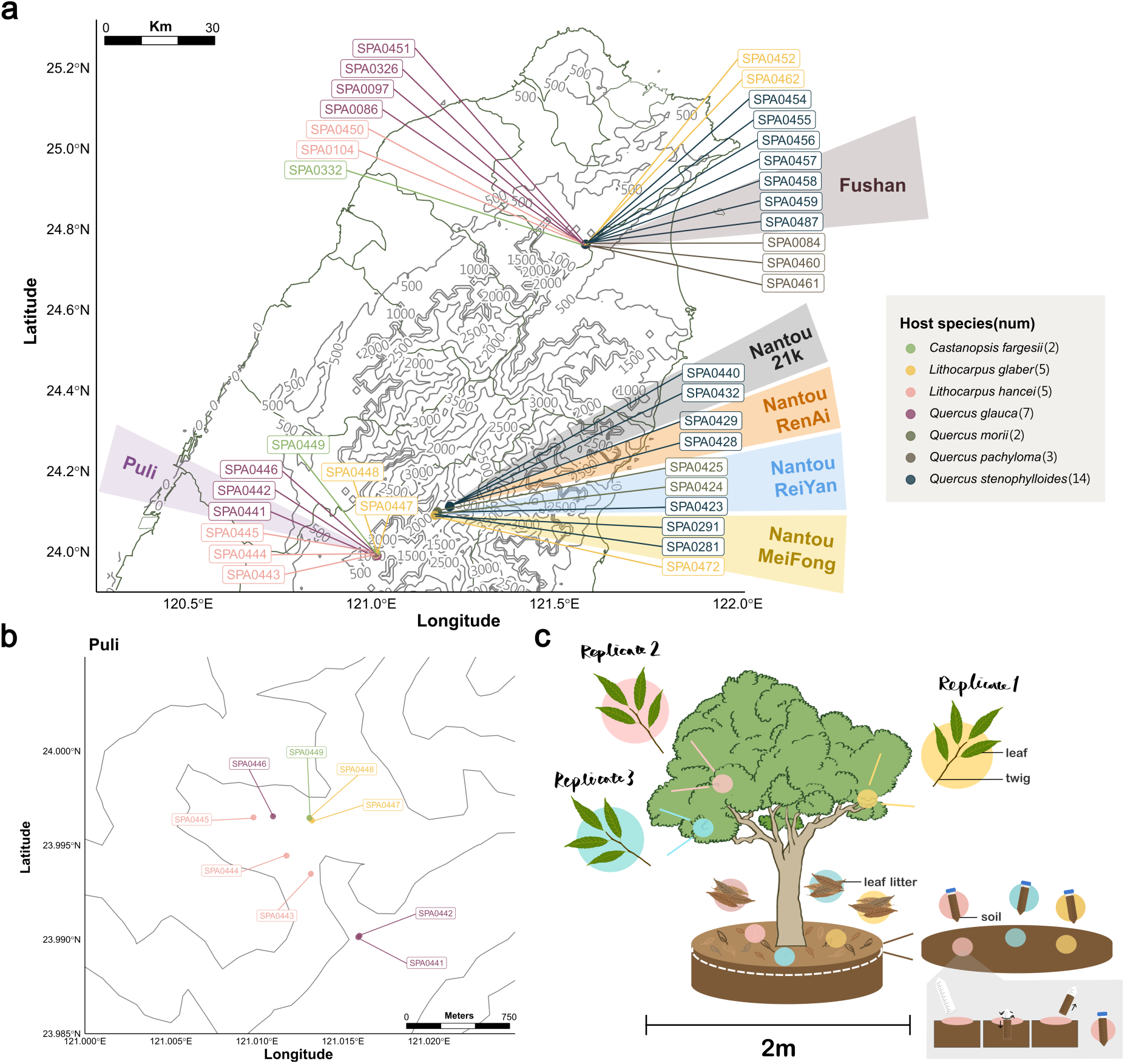
Sampled broadleaf forests in Taiwan surveyed for fungi associated with Fagaceae species. (a) Map of sampling sites. (b) Tree distributions in the Nantou Puli site. (c) A schematic diagram showing leaf, twig, leaf litter, and soil were sampled triplicates in each tree.

For each tree, triplicate samples of each habitat were obtained from different sides of the tree (**Fig. 1c**). The twigs, comprising approximately 20-30 leaves, were pruned with a sterilised tree lopper and placed into seal bags, with the leaves and twigs separated during laboratory preprocessing. Leaf litter (comprising 20-45 leaves) was sampled from the organic layer of soil, consisting mostly of late-stage decomposition material [30], while topsoil was sampled at a depth of 0–10 cm using 50 ml falcon tubes after the organic layer was removed. Originally 18 trees were chosen per survey site (a total of 36), but due to decay, two additional trees were substituted (SPA0446 and SPA0457 were replaced with SP0472 and SPA0487, respectively). A total of 864 samples were collected. All samples were refrigerated at 4°C until extraction of genomic DNA (**Supplementary Table S1**). Abiotic variables associated with survey sites were collected from Climate Observation Data Inquire Service, Taiwan (**Supplementary Table S2;** https://codis.cwa.gov.tw/).

### Sample preprocessing and DNA extraction

The preprocessing steps for leaf, twig, and leaf litter samples followed the protocols detailed in our previous study [25]. Genomic DNA was collected from sample surfaces using 0.22 μm PES membranes in filtration cup (Jet Bio-Filtration Co., Cat. FPE214250). As a control for background noise, three sterilized filter papers were processed in parallel. The genomic DNA was then extracted using the DNeasy PowerWater kit (QIAGEN; Cat. 14900-50-NF) following the manufacturer’s instructions.

The topsoil samples were sieved using 2 mm steel mesh to remove plant debris, insects and rocks. Genomic DNA was extracted from approximately 0.25 g of soil using the DNeasy PowerSoil Pro kit (QIAGEN, Cat. 47014) as instructed by the manufacturer. Homogenisation was performed with a Precellys 24 Touch homogeniser (Bertin Technologies, Cat. P002391-P24T0-A.0) at 5000 rpm for two cycles of 90 seconds, with a 15-second pause between cycles. The extracted DNA was quantified with Invitrogen Qubit 4 fluorometer (Invitrogen) and NanoDrop 1000 (ThermoFisher) and stored at −20°C until library preparation.

### Amplicon library construction and sequencing

Amplicon libraries were constructed as previously described [31] using the forward primer ITS3ngs (5-CANCGATGAAGAACGYRG-3’) and reverse primer ITS4ngsUni (5’-CCTSCSCTTANTDATATGC-3’) [32, 33] to amplify the ITS2 region. The PCR mixture contained 50 ng of DNA extract, 2 μl of each 10 μM primer, 8 μl 5x HOT FIREPol Blend Master Mix (Solis Biodyne, Cat. 04-27-00115), 1 μl of 25 mM MgCl_2_ and ddH_2_0 to 40 μl. Extracted genomic DNA of *Saccharomyces kudriavzevii*, *S. paradoxus* and *S. cerevisiae* were used as positive controls to assess the false positives in the sequencing and analysis. The thermal cycling conditions consisted of an initial denaturation at 95°C for 12 min, followed by 35 cycles of denaturation at 95° C for 20 sec, annealing at 55°C for 30 sec, and extension at 72°C for 1 min, finishing with a final cycle at 72°C for 7 min. Biological replicates planned to be merged and sequenced were pooled with equal molar. Amplicons were normalised to equal DNA quantity (approximately 25 ng) using SequalPrep^TM^ Normalization Plate Kit (Invitrogen, ID: A1051001) according to the manufacturer’s instructions before pooling. The pooled library was concentrated to 10 ng/μl using AMPure XP (Beckman Coulter, ID: A63881). Each batch produced two plates of the library. Libraries were sequenced using the Illumina Miseq PE300 sequencing platform with equal molar pooling and 20% Phix spike-in.

### Statistical analyses

The raw sequencing data were imported and demultiplexed using *sabre* (v1.0; https://github.com/najoshi/sabre) allowing a 1 bp mismatch. Sequencing quality was examined using *FastQC* (v0.11.9; https://github.com/s-andrews/FastQC). Reads without primer sequences were detected and discarded with *usearch* (v11.0.667) [34]. Primer sequences were trimmed using *Cutadapt* (v4.4) [35]. The filtered and trimmed sequences were processed with the *Qiime2* (v2023.5.1) [36] pipeline to filter reads with a quality threshold of Qscore > 20 and to denoise into amplicon sequence variants (ASVs). Taxonomy for the ASVs was assigned using *constax* (v2.0.18) [37] with the UNITE Fungal database (v9.0) [38], and the fungal guild was annotated using *FUNGuild* (v1.2) [39].

Data processing and analysing as following were performed in the R-studio environment (v2024.4.1.748) [40]. The taxonomy levels were updated using package *rgbif* (v3.7.7) [41]. Background reads were subtracted based on the median read number in negative controls, and ASVs with relative abundances below 0.1% in each sample were filtered out, informed by positive control results. Preprocessed sequencing data were analyzed with *phyloseq* (v1.40.0) [42]. Figures were generated using *ggplot2* (v3.4.2) [43]. The sampling locations were annotated the sampling locations using *ggspatial* (v1.1.9; https://CRAN.R-project.org/package=ggspatial), *metR* (v0.14.0; https://github.com/eliocamp/metR) and *ggrepel* (v0.9.3; https://CRAN.R-project.org/package=ggrepel). Statistical significance test of the alpha diversity index was performed using *HSD.test* function in *agricolae* package (v1.3.7; https://github.com/myaseen208/agricolae). ASVs present in at least 25% of samples within a single habitat were classified as endemic, while those occurring in 25% of samples across multiple habitats (e.g., 25% in both soil and leaf samples) were designated as ubiquitous ASVs. The *UpSetR* package was performed to visualise the endemic and ubiquitous ASVs through the environments (v1.4.0; https://CRAN.R-project.org/package=UpSetR).

The normalised stochasticity ratio (NST) based on Jaccard distance was calculated to assess mycobiome assembly processes using the *NST* package (v3.1.10) [44]. A random forest model was used to classify samples into their respective habitats and to analyse ASV importance based on the mycobiome composition using package *randomForest*(v4.7.1.1; https://github.com/cran/randomForest/). Optimised parameters, including ntree, train/test ratio, mtry and minimum node size, were tuned using package *ranger* (v0.16.0; optimised parameters were as followed: mtry=70; minimum node size=1; ntree=3000) [45]. Samples with fewer than 5,000 reads were excluded from the analysis. A total of 679 samples were initially transformed to relative abundance and subsequently divided into training set and testing set with an 80/20 ratio using function createDataPartition from package *caret* (v6.0.94) [46]. Cross-validation was used to prevent model overfitting, and model discrimination was visualized with the *pROC* (v1.18.5) [47] package.

Sample heterogeneity was calculated with the vegdist function from the *vegan* package (v2.6-4; https://CRAN.R-project.org/package=vegan), focusing on samples with triplicates that underwent separate sequencing. Samples with fewer than 5,000 reads were excluded before analysis, and ASV relative abundance was normalised by square-rooting transformation. Comparisons were divided into three categories: ‘Within tree’ indicated the comparison between biological replicates of the same tree, ‘Within site’ referred to comparing samples from different trees in the same sampling sites and season and ‘Between sites’ showed the comparison of samples from different trees and sites in the same season (Spring of Puli vs. Spring of Nantou; Fall of Puli vs. Fall of Nantou).

### Network analysis

Leaf, twig, and litter data were rarefied to 15,000 reads per sample, while soil data were rarefied to 10,000 reads per sample. ASVs present in more than 50% of the habitat samples were retained by applying a 50% prevalence threshold to the rarefied data. Correlation indices were calculated using *FastSpar* (v1.0.0) [48, 49] with 100 iterations and filtered based on both significance (false positive adjusted p-value < 0.05) and strength (SparCC ≥ 0.6 or SparCC ≤ −0.6). The co-occurrence networks were visualised using *igraph* [50]. To identify putative keystone taxa, we estimated the Zi (within-module degree z-score) and Pi (participation coefficient) values for each node using function within_module_deg_z_score and part_coeff of *brainGraph* (v3.0.0; https://CRAN.R-project.org/package=brainGraph) package, respectively. Nodes with Zi greater than 2.5 were defined as modular hubs; those with Pi more than 0.62 were identified as connectors; and nodes that satisfied both conditions of Zi ≥ 2.5 and Pi ≥ 0.62 were classified as network hubs.

## Results

### Fungal diversity across habitats and seasons in Taiwanese Fagaceae forests

We surveyed 38 trees from seven Fagaceae species across two natural Taiwanese broadleaf forests and Fushan botanical garden (**Fig. 1**). The survey was conducted at two time points and encompassed four habitats: leaf, twig, leaf litter, and topsoil, with triplicates for each tree resulting in a total of 864 samples (**Fig. 1c**). By amplifying and sequencing the ITS2 region, we quantified the relative abundances of mycobiomes across microhabitats and seasons. After quality filtering, denoising, merging and removing false positives, we obtained 17,222,990 sequences from an initial 46,317,215 paired reads and classified them into 11,600 amplicon sequencing variants (ASVs), averaging 69 ASVs and a median relative abundance of 0.40% per sample.

The ASVs were classified into seven phyla with 14.4% remaining unclassified. The dominant fungal phylum was Ascomycota (55.62%), followed by Basidiomycota (26.6%), Zygomycota (10.46%), Glomeromycota (0.40%), Chytridiomycota (0.39%), and Blastocladiomycota (0.02%) (**Supplementary Fig. S1**). ASV1 was identified belonging to the *Cladosporium* genus and was the most dominant ASV present in 332 out of 736 samples with an average relative abundance of 5.05% (**Fig. 2a**). This ASV was particularly prevalent in leaf litter and leaves, with average relative abundance of 6.21% and 4.89%, respectively, in comparison to 0.008% for other ASVs.

**Fig. 2.**
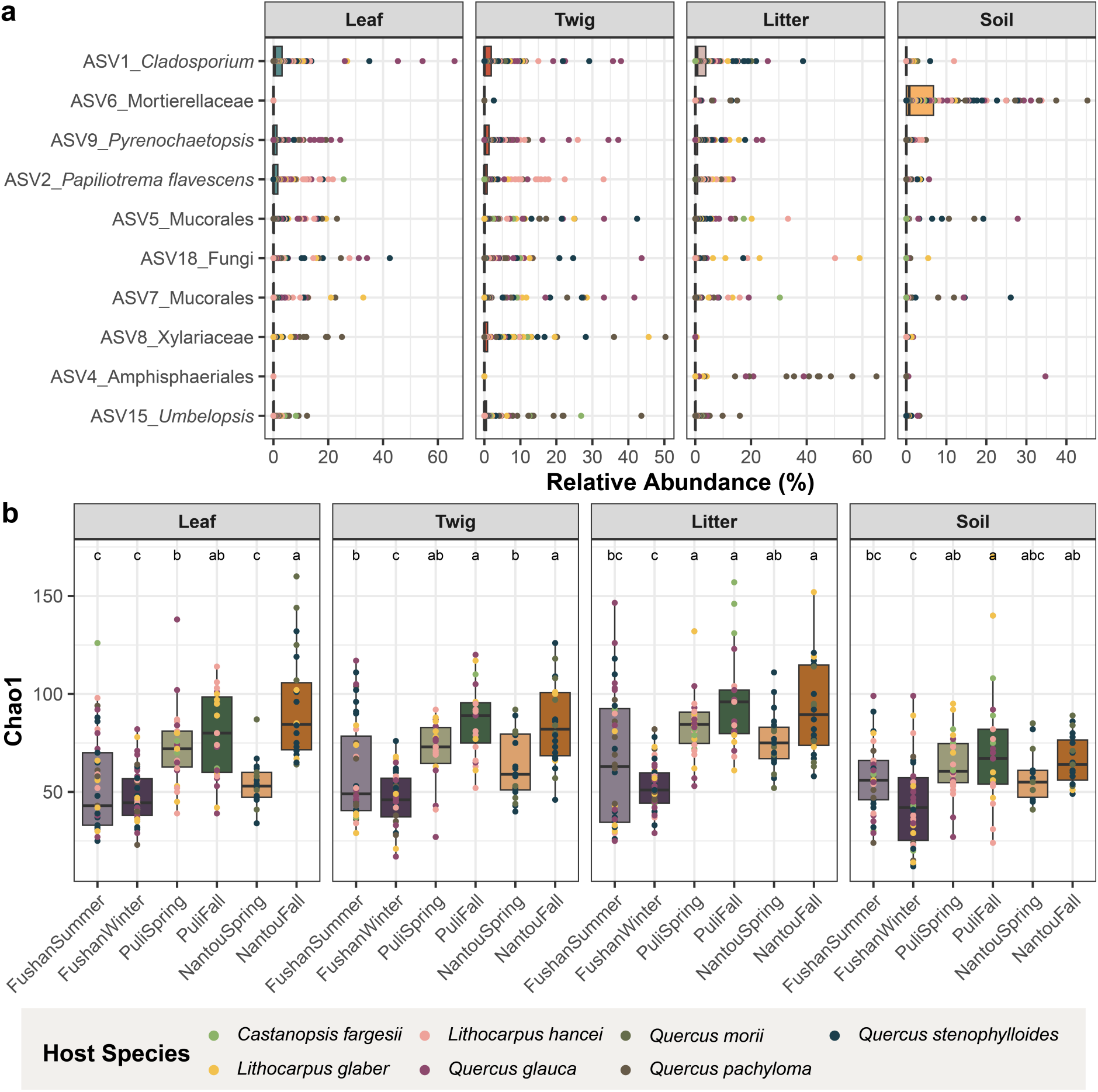
Fungal community diversity. (a) Top 10 ASVs’ relative abundance across samples. Each dot represents one sample, the color indicates the host species and the shape denotes the collection seasons; (b) Alpha diversity across habitats and seasons calculated using Chao1 index. The significant difference test was performed using Tukey’s HSD test, with different letters indicating statistically significant differences (P < 0.05). Dot color represents host species.

Alpha diversity, assessed using the Chao1 and Shannon diversity indices, revealed significant variation across habitats, with higher richness and evenness in litter (73 ± 27 ASVs) compared to twig (65 ± 24 ASVs) and leaf (64 ± 26 ASVs) (**Fig. 2b** and **Supplementary Fig. S2**), while soil exhibited the lowest diversity (58 ± 22 ASVs). Seasonal variation also emerged as another significant driver of fungal communities (PERMANOVA; R^2^ = 0.09, P < 0.001). For instance, at the Fushan Botanical Garden, fungal richness and evenness declined in winter compared to summer, while in Nantou and Puli, richness increased in fall compared to spring, though evenness remained stable (**Fig. 2b** and **Supplementary Fig. S2a**). There were no significant differences in both Chao1 and Shannon indices at the host species level, suggesting that host identity at a genus level may not play a major in determining fungal diversity in Fagaceae broadleaf forests. An exception occurred in Puli during the fall, where alpha diversity varied among host species, with *Castanopsis fargesii* exhibiting the highest diversity (**Supplementary Fig. S2b**). These findings suggest that while habitats and season are key determinants of fungal community diversity, host species may exert a more localized or context-dependent influence on alpha diversity. (**Supplementary Fig. S2b**).

### Spatiotemporal variation in fungal diversity and niche assembly of broadleaf forests

To investigate the spatial heterogeneity of fungal communities, we calculated the Bray-Curtis (BC) dissimilarity index based on the square root-transformed relative abundance of samples collected from the same trees, within less than 1 meter apart (**Fig. 1c and Fig. 3a**). Fungal community from the same microhabitat showed high dissimilar even within individual trees (median BC = 0.66, **Fig. 3a**). Samples originating from the same habitats were more similar than those between different habitats (median BC = 0.66 vs. 0.94, **Fig. 3a**). Further partitioning of the BC dissimilarity revealed differences were driven by regions. For example, two *Quercus morii* and one *Q. stenophylloides* trees located in Rueiyan of the forest in Nantou exhibited the lowest BC values (**Supplementary Fig. S3**). Among habitats, leaves exhibited the lowest dissimilarity across most seasons, suggesting that leaves may support a more stable mycobiome composition within the same tree (**Fig. 3a and Supplementary Fig. S4**). Leaf versus twig comparisons had BC values similar to intra-habitat comparisons, indicating that fungal communities in these two habitats were alike (**Fig. 3a and Supplementary Fig. S5**). Overlap between litter vs. soil and litter vs. leaf, i.e., exhibiting low BC values, was also observed, likely due to fungal movement between microhabitats.

**Fig. 3.**
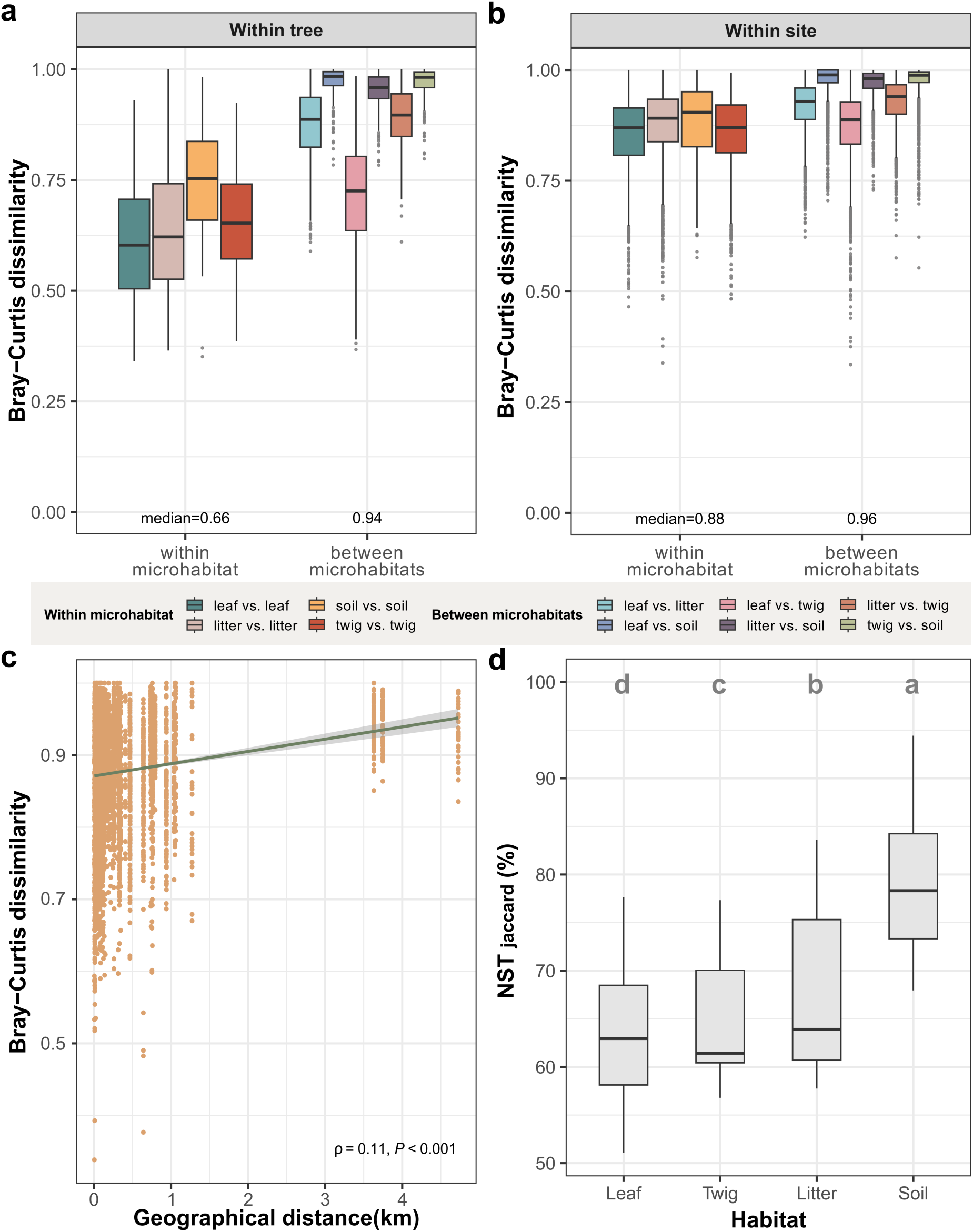
Dissimilarity and ecological stochasticity in the fungal community associated with Fagaceae species across habitats and geographical distance (km). Bray-Curtis dissimilarity of samples across microhabitats from the same tree (a) and different trees in one site (b). The colours indicate the microhabitat pairwise of two samples. (c) Distance-decay relationship of sample similarity of the same microhabitat between tree host species. Spearman’s rank correlation coefficient (ρ) and the corresponding significance are provided to demonstrate the strength and significance of the distance-decay relationship. (d) Community assembly processes are calculated by normalised stochasticity ratio (NST) based on Jaccard distance among habitats. The letters represent the statistically significant differences as estimated by Tukey’s HSD test (P < 0.05).

When considering distances between trees from the same forest site, samples from the same habitats remained highly dissimilar (median BC = 0.88; **Fig. 3b**) and a clear distance-decay relationship was observed (**Fig. 3c**). This relationship was stronger among trees of the same species compared to those of different species (Spearman’s ρ = 0.42 vs 0.11, respectively; **Fig. 3c and Supplementary Fig. S6**), suggesting that host species played a significant role in determining fungal community composition at shorter distances, whereas communities between distantly located trees will be highly dissimilar, even if they belong to the same species. The distance-decay relationship was most pronounced in leaf vs. leaf comparisons, especially in Nantou during the Fall and Spring seasons (Spearman’s ρ = 0.54 and 0.44, respectively, **Supplementary Fig. S7**). In contrast, soil vs. soil comparisons displayed weaker or insignificant distance-decay relationships. As expected, samples from different microhabitats, trees and sites exhibited the greatest dissimilarity (median BC = 0.99; **Supplementary Fig S8**), suggesting that distinct habitats act as strong barriers to fungal dispersal regardless of physical proximity. In addition, we analysed the normalised stochasticity ratio (NST) to evaluate the balance between stochastic and deterministic processes in fungal community assembly across different habitats and seasons (**Fig. 3d**). Leaves exhibited the lowest stochasticity (mean NST = 0.64), followed by twigs and litter, while soil had the highest stochasticity (mean NST = 0.79).

### Distribution, functional guilds, and predictors of ASVs across forest habitats

To characterise the distributions of ASVs, we designated their status as endemic and ubiquitous based on their presence across different habitats (see Methods; **Fig. 4a**) and annotated their functional guilds against the FUNGuild database [39]. Soil harbored the highest number of endemic ASVs, with the saprotroph-symbiotroph functional guild being the most prevalent, showing an average relative abundance of 3.43% across all soil samples (**Fig. 4b**). Despite the close physical proximity between soil and litter, no ASVs were shared between these two habitats under our filtering criteria. Only two ASVs-*Cladosporium* (ASV1) and *Pyrenochaetopsis* (ASV9)-were identified as ubiquitous across all habitats. In contrast, the above-ground habitats, encompassing leaves, twigs, and litter, shared 12 ASVs, which collectively account for approximately 8.89% of the relative abundance in each sample (**Fig. 4b; Supplementary Fig. S9**). Pathogenic fungi were predominantly found in above-ground habitats, with leaves exhibiting the highest abundances.

**Fig. 4.**
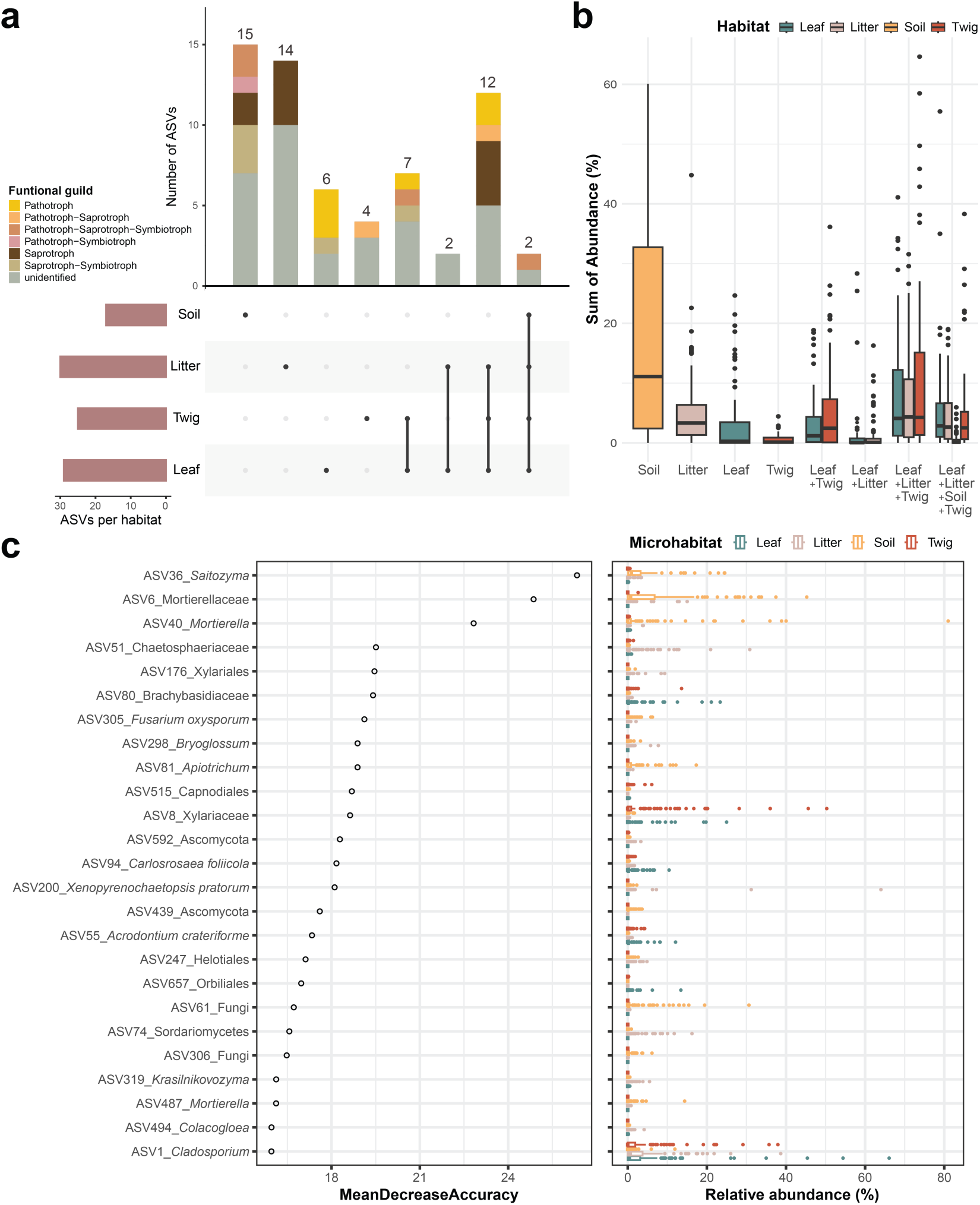
The endemicity of ASVs and their role in the random forest model. Number (a) and relative abundance (b) of endemic and ubiquitous ASVs among habitats. Habitat combinations without sharing ASVs are not shown. (a) The bar color indicates the guild of the ASVs; (b) Each dot represents the sum of the endemic/ubiquitous ASV abundances in one sample. The box color represents the compartment. (c) Importance of ASVs in microhabitats classification and their relative abundance. The colors of boxes and dots represent its microhabitat.

We employed random forest analysis to classify habitats based on ASV abundances and to assess the importance of specific ASVs in distinguishing between habitats (**Fig. 4c**). The model demonstrated strong predictive power, with an out-of-bag (OOB) error rate of 9.67% on training data and 11.02% on validation data (mean AUC = 0.982; **Supplementary Fig. S10 and Supplementary Table S3a**). Misclassification primarily occurred between leaves and twigs, with 31 out of 136 twigs misclassified as leaves (OOB error rate = 22.79%) and 11 out of 141 leaves misclassified as twigs (OOB error rate = 7.8%), consistent with the general low BC values in samples between these two habitats. Some litter and soil samples were classified incorrectly as leaf or twig, but not vice versa. Specifically, 7 out of 142 litter samples were misclassified as leaf and 5 out of 127 soil samples were misclassified into other three habitats. Merging leaf and twig into a single category reduced the OOB error rate to 6.96%, with 100% accuracy for the combined category (**Supplementary Table S3b**), suggesting vertical movement of fungal communities from canopy to ground.

Several ASVs emerged as strong predictors of specific habitats. For example, ASV36 (*Salilomyza*) and ASV6 (assigned to the family Mortierellaceae) were key predictors for soil with relatively high relative abundances in soil samples (**Fig. 4c**), while ASVs such as ASV80 (assigned to the family Branchybasidiaceae) were more abundant in leaf samples, highlighting their role in shaping the unique fungal communities in the phyllosphere.

### Environmental predictors of factors on forest mycobiome

Fungal composition differences were analysed using Bray-Curtis distance and visualised through Non-metric Multidimensional Scaling (NMDS) and Principal Coordinates Analysis (PCoA), with the first two principal components in PCoA explaining 13% of the variation in the dataset (**Fig. 5 and Supplementary Fig. S11**). Both NMDS and PCoA analyses revealed that samples collected from the same location but during different seasons significantly cluster together (ANOSIM R = 0.47, P < 0.001). Among habitats, NMDS analysis showed clear distinctions between soil and the other three habitats (leaf, litter, twig) (ANOSIM R = 0.31, P < 0.001) (**Fig. 5a**). Although host species had a significant influence on the fungal composition, the effect was relatively minor (ANOSIM R = 0.18, P < 0.001). Despite similar altitudes between the Fushan botanical garden (625-660 m) and the Puli forest (593-774 m), samples from these sites clustered separately.

**Fig. 5.**
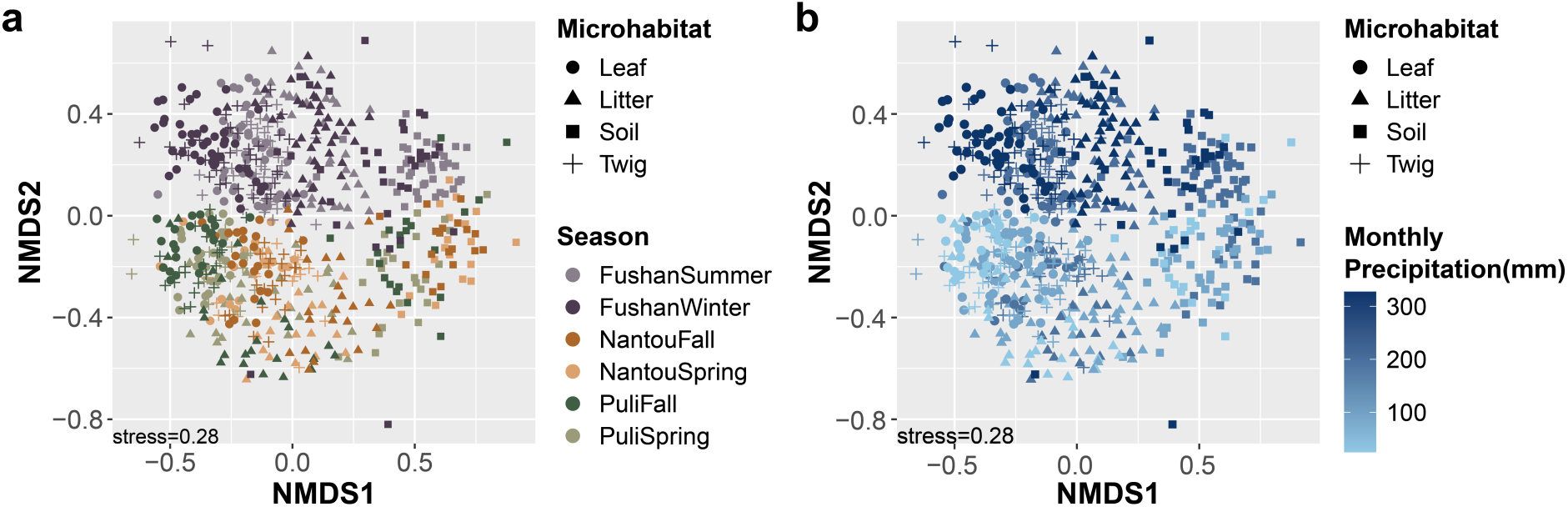
Fungal community differences among habitats, seasons and monthly precipitation. The plots were calculated using the Bray-Curtis distance, ordinated using Non-metric Multidimensional Scaling (NMDS) analysis. Samples with less than 5,000 reads were discarded before analysing. This analysis was performed at the genus level. (a) The shapes represent the microhabitat of the sample. The colors indicate the season of the samples; (b) The shapes represent the microhabitats and the colors represent the monthly precipitation (mm).

Given the significant role of seasonality in shaping mycobiome composition, we explored the influence of climatic variables, focusing on relative humidity, temperature, and precipitation. Among these, monthly precipitation emerged as the most influential in determining mycobiome composition (PERMANOVA R² = 0.016, P < 0.001). PCoA based on Bray-Curtis distance, showed that PC1 (8.1%) effectively separated soil samples from the other habitats, while PC2 (4.9%) distinguished samples based on monthly precipitation levels, with higher precipitation associated with distinct fungal communities (**Supplementary Fig. S11b**). In contrast, daily precipitation had a weaker effect on fungal composition (PERMANOVA, R² = 0.007, P < 0.001), suggesting that prolonged rainfall, rather than short-term precipitation, plays a more critical role in gradually influencing fungal community composition.

### Complexity and connectivity of co-occurrence networks across habitats

The co-occurrence network was constructed from 1,313,817 sequences representing 517 prevalent ASVs, revealing how different habitats and seasons influence fungal complexity and connectivity (**Fig. 6; Supplementary Fig. S12**). Significant variations in network size and complexity across different habitats and environmental conditions were observed. Soil networks were notably smaller and less complex than those of the other three habitats, with an average of 14 ASVs compared to 39 ASVs in leaves, litters, and twigs. This finding underscores the distinct ecological dynamics within the soil, which may be more isolated or less interconnected than other habitats. Interestingly, the Fushan botanical garden exhibited a significantly lower network size and complexity than the other two natural forests, suggesting that the human-managed environment results in reduced fungal-fungal interactions compared to more natural environments. Seasonal comparisons revealed that network complexity decreased in winter compared to summer, with increases in the leaf of puli, twig of puli, litter, and soil, and decreases in the leaf and twig of Nantou from spring to fall **(Supplementary Fig. S12)**, indicating that seasonal shifts, including changes in temperature and humidity, significantly impact fungal community interactions. Modularity, reflecting how a network divides into distinct modules, varied significantly across habitats and seasons (**Fig. 6 and Supplementary Fig. S12**). For instance, in Fushan, the modularity of leaf and litter networks increased from summer to winter (modularity of summer vs. winter: leaf = 0.59 vs. 0.67, litter = 0.58 vs. 0.75), while twig and soil modularity decreased (modularity of summer vs. winter: twig = 0.70 vs. 0.52, soil = 0.51 vs. 0.22). Generally, the modularity and network complexity increased from spring to fall in leaves but decreased in litter, with twig and soil networks remaining relatively stable.

**Fig. 6.**
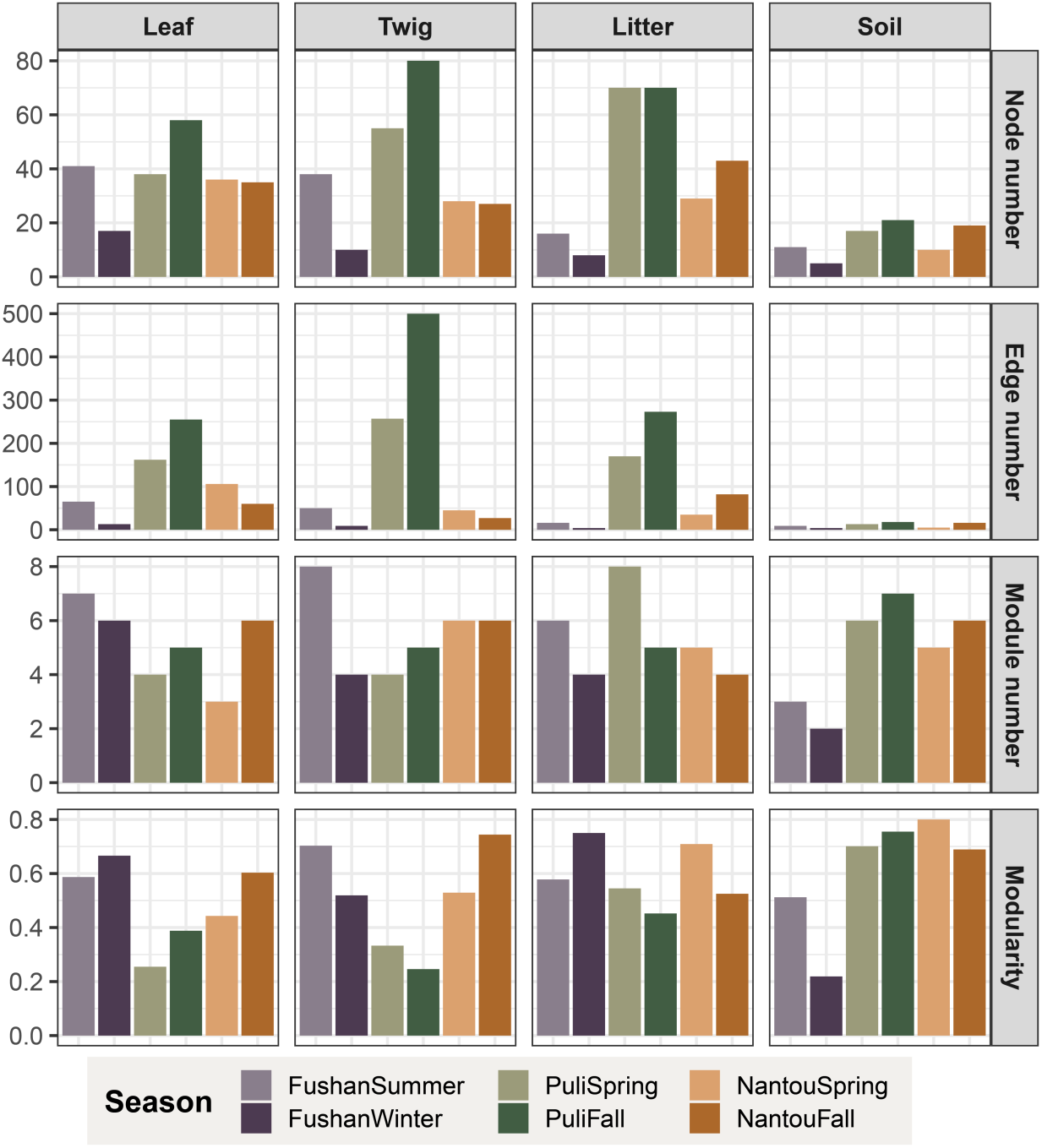
Complexity and connectivity of co-occurrence networks across seasons and substrates demonstrated by node number, edge number, module number and modularity.

We found significant negative correlations between NST and both node and edge numbers (Spearman’s ρ= −0.65, P < 0.001 for nodes; Spearman’s ρ= −0.71, P < 0.001 for edges; **Supplementary Fig. S13**), indicating that as the number of nodes (ASVs) and edges (interactions) within a network increase, the influence of stochastic processes in community assembly decreases. Conversely, no significant correlations were observed between NST and the number of modules or modularity. The proportion of functional guild nodes also varied across seasons, with an increase in the proportion of saprotroph nodes in soil and saprotroph and pathotroph-saprotroph nodes in litter during the fall (**Supplementary Fig. S14**), highlighting the role of environmental changes in shaping fungal community composition.

To assess the importance of individual nodes, we examined their connectivity within (modular hubs: Zi ≥ 2.5) and between (connectors: Pi ≥ 0.62) modules (see Methods; **Supplementary Fig. S15**), designating them as modular hubs or connectors based on criteria set in [51]. No shared modular hubs or connectors were found between seasons, even within the same location, and none were identified in soil or during winter. In leaf networks, eight connectors were identified, including two pathotroph-saprotroph-symbiotroph connectors (ASV9_*Pyrenochaetopsis*, Pi = 0.67, and ASV102_*Cryptococcus*, Pi = 0.65) and six unidentified connectors (**Supplementary Table S4**). The twig network had two connectors, while the litter network exhibited nine connectors (**Supplementary Table S4**) and one modular hub (pathotroph ASV614_*Dactylaria* acacia, Zi = 2.57). Only one putative keystone species—ASV1_*Cladosporium* (Pi = 0.66, Zi = 2.87)—was recognized as a network hub in the leaf of Puli during spring. Connectors were more frequently identified in leaves, twigs, and litter in Puli across both spring and fall, while in Nantou and Fushan, connectors were primarily observed in leaves or litter. Together these patterns suggest that warmer and drier climates may lead to lower modularity, larger modules, and increased opportunities for observing connectors within fungal networks.

## Discussion

This study offers a detailed quantification of the spatiotemporal heterogeneity in fungal community composition within various above-ground habitats in tropical Fagaceae forests. By examining fungal communities in leaves, twigs, litter, and soil, we provide comprehensive insights into how different microhabitats influence fungal diversity at a local scale. We revealed that the alpha diversity in the phyllosphere and litter fungi was higher than that observed in soil fungi, a pattern that contrasts with observations in bacterial microbiomes, where alpha diversity is typically highest in soil and lowest in the forest foreground [52]. This highlights the complexity of fungal community dynamics and underscores the necessity of considering multiple habitats to fully understand broader ecological processes.

At small spatial scales, high BC dissimilarity (median 0.66) was observed among fungal communities from the same habitat, even when samples were collected from the same tree, consistent with previous studies that reported highly dissimilar fungal communities in litter fungi over short distances [21]. The level of dissimilarity varied significantly between microhabitats, with leaves exhibiting the lowest dissimilarity across seasons, suggesting a relatively stable mycobiome composition. In contrast, litter and soil showed higher variability, likely due to their dynamic nature and exposure to fluctuating environmental conditions. Our analysis of the normalised stochasticity ratio (NST) further supports these observations, with leaves showing lower NST values, indicating a stronger influence of deterministic processes, while litter and soil exhibited higher NST values, reflecting greater stochasticity in these more dynamic habitats. At this small spatial scale, BC dissimilarity almost plateaued between fungal communities (median 0.94), suggesting a strong biogeography barrier resulting in disjunctive fungal communities. The only exception was leaf vs. twig, resembling intra-habitat differences. The highest inter-habitat differences were observed in the leaf vs. soil and twig vs. soil, reflecting the substantial heterogeneity in forest ecosystems, where physical separation and environmental factors collectively drive fungal diversity.

Our research uncovered distinct spatial distance-decay patterns in fungal communities across various habitats. Specifically, the leaf, twig and litter fungi displayed relatively low baseline community similarity and distance-decay relationships were largely influenced by site and season. For example, in the Nantou region during spring, litter samples displayed a high correlation (Spearman’s ρ = 0.74), while no significant correlation was found in Puli during the same season. This suggests that while these fungi can disperse over short distances, environmental selection exerts a crucial influence on community assembly [11]. In contrast, soil fungal communities exhibited no or weak distance-decay relationships, which aligns with previous studies that found weaker distance decay in soil compared to other habitats [13, 53]. These findings underscore the role of microhabitat-specific factors in shaping fungal community dynamics and emphasize the need to consider these factors when studying fungal biogeography in forest ecosystems.

Documenting fungal communities across multiple microhabitats at small spatial scales allowed us to identify instances of overlap, with approximately 6% of samples from one microhabitat being more similar to samples from another microhabitat. This overlap suggests potential pathways for fungal movement or shared environmental conditions that facilitate community convergence and reinforce the importance of spatial orientation and environmental context in the dispersal and establishment of fungal communities. For example, certain litter samples showed similarities with soil communities, likely due to the physical proximity and interaction between these habitats [21]. This pattern emphasizes the importance of microbial-mediated interfaces, such as the litter-soil interface, in shaping community composition [20]. Moreover, the presence of similar communities across distinct habitats highlights the role of dispersal processes and environmental filtering in shaping fungal communities [11]. Dispersal limitation can lead to greater community similarity across neighboring habitats, as demonstrated by studies where fungal communities in leaves and twigs were more similar than expected due to their connectedness through dispersal vectors [54]. This indicates that while microhabitats can support distinct communities, interactions at their interfaces can lead to the blending of fungal assemblages at short timescales. Interestingly, such interactions can lead to misclassification of sample origin in the random forest analysis (error rate 9.68%), and we observed no misclassification in leaf and twig samples as litter or soil, possibly due to the reduced likelihood of upward transmission compared to horizontal movement across other substrates.

On a broader scale, environmental factors like seasonal changes and long-term precipitation (abiotic effects) exert a more significant role [11, 55]. Among climatic variables, monthly precipitation had the most pronounced impact on fungal diversity, influencing both mycobiome composition and richness. The NST analysis further demonstrated that microhabitats with higher complexity and connectivity, such as leaf and twig, were primarily influenced by deterministic processes, whereas more isolated or less complex habitats like soil were governed by stochasticity. This scale-dependent variation is consistent across different habitats, with leaves showing particularly stable fungal communities over time [56], whereas soil shows the most fluctuations. Our comprehensive dataset, encompassing various spatial scales and environmental conditions, highlights the dynamic and complex nature of fungal biogeography [9]. Contrary to the previous study, the host identity in our study did not have a discernible impact on either alpha diversity or fungal composition [57]. However, the results of beta diversity analyses indicated that fungal communities in samples from higher altitudes, such as Nantou 21k, Nantou RenAi, Nantou ReiYan, and Nantou Meifong, were more similar to each other than to those from lower altitudes like Fushan and Puli. This suggests that altitude strongly influences fungal composition, following Rapoport’s rule, which posits that species in environments with greater variability in conditions are expected to have larger ecological tolerances and, consequently, broader ranges [58].

An unidentified *Cladosporium* (ASV1) serves as the sole network hub in the co-occurrence network analysis and a key predictor on random forest analysis (mean decrease accuracy = 15.96). ASV1 also emerged as the most dominant species in the study, particularly in leaf and litter samples, as indicated by studies that have revealed *Cladosporium cladosporioides* as a common species in the phyllosphere and on fallen leaves [59]. A subsequent investigation of the *Cladosporium* strain further substantiated its capacity to degrade various forms of humus, including laccase, lignin and triarylmethane [60]. The prevalence of *Cladosporium* in these habitats highlights its potential role in leaf composition, nutrient cycling and organic matter decomposition, thereby emphasizing its ecological importance in phyllosphere fungal communities. In addition, *Saitozyma* (ASV36) and *Apiotrichum* (ASV81) were important predictors for habitat classification and were predominantly found in soil. This dominance is supported by prior findings from European Fagaceae forests [61], underscoring broader ecological importance beyond local habitats. *Apiotrichum* species have been demonstrated to degrade phenolic compounds [62, 63], hemicelluloses [64] and benzene compounds [65], while *Saitozyma* species possess the capacity for uptake of carbon from cellulose [66], suggesting their role in the decomposition of plant biomass. These findings provide valuable insights into the assembly processes of fungal communities and illustrate the pivotal role of the fungal ecological function in shaping ecosystem dynamics.

## Conclusion

By examining spatiotemporal heterogeneity across coexisting habitats, we reveal contrasting processes driving fungal community assembly in tropical broadleaf forests. Deterministic influences dominate in the interconnected leaf and twig habitats, while stochasticity is more prominent in soil and litter, highlighting the role of habitat features in shaping communities. The identification of *Cladosporium* sp. as a keystone species further highlights the ecological importance of certain taxa in maintaining network stability and community cohesion. Random forest analysis identified distinct fungal signatures, with misclassification linked to natural overlap between habitats. Our findings emphasize the importance of multi-habitat approaches for understanding fungal biogeography and underscore the need to explore habitat-specific drivers for ecosystem functioning and conservation in response to climate change.

## Data availability

The demultiplexed Illumina data were deposited to NCBI under BioProject PRJNA1156365. The accession number of each sample were complemented in **Supplementary Table S1**.

## Acknowledgements

We would like to thank Chen Hsiao, Daphne Z. Hoh, Wei-Ting Chien and Chia-Chi Huang for assistance during sampling.

## Author information

### Contributions

IJT conceived and led the study. CPL and YCL carried out the sampling. CPL and YFL conducted the metabarcoding experiments and CPL performed amplicon data analysis with guidance from YFL. HMK performed the laboratory work. MYJL performed the sequencing. CPL and IJT wrote the manuscript. All authors reviewed and approved the final version of the manuscript.

## Ethics declarations

### Ethics approval and consent to participate

Not applicable.

### Consent for publication

Not applicable.

